# Sex dependent gene activity in the human body

**DOI:** 10.1101/2020.04.22.055053

**Authors:** Robin J.G. Hartman, Michal Mokry, Gerard Pasterkamp, Hester M. den Ruijter

## Abstract

Many pathophysiological mechanisms in human health and disease are dependent on sex. Systems biology approaches are successfully used to decipher human disease etiology, yet the effect of sex on gene network biology is mostly unknown. To address this, we used RNA-sequencing data of over 700 individuals spanning 24 tissues from the Genotype-Tissue Expression project to generate a whole-body gene activity map and quantified the sex differences per tissue. We found that of the 13,787 genes analyzed in 24 tissues, 20.1% of the gene activity is influenced by sex. For example, skeletal muscle was predominantly enriched with genes more active in males, whereas thyroid primarily contained genes more active in females. This was accompanied by consistent sex differences in pathway activity, including hypoxia, epithelial-to-mesenchymal transition, and inflammation over the human body. Furthermore, multi-organ analyses revealed consistent sex-dependent gene activity over numerous tissues which was accompanied by enrichment of transcription factor binding motifs in the promoters of these genes. Finally, we show that many sex-biased genes are known druggable targets. This emphasizes sex as a biological variable and the need to incorporate sex in systems biology studies.

## Introduction

Sex influences the processes underlying development and disease. Sex differences in the prevalence of diseases are appreciated nowadays, e.g. autoimmune diseases are more common in females, whereas non-reproductive cancers are more common in males^1^. At the basis of this diversity in disease prevalence are the molecular and genetic differences between male and female cells driven by sex steroids and sex chromosomes^2^. Previous efforts have shown that the human transcriptome is sex-biased over different tissues^3^.

Systems and network biology are successfully used to find disease targets based on the complexity of biological systems, and recent studies have shown that sex is involved in shaping biological networks^4,5^. However, a whole-body map of sex differences in gene activity is lacking, while gene activity underlies biological gene networks. We hypothesize that substantial and overarching sex differences exist in gene network activity in tissues of the human body.

Therefore, we created a whole-body map of sex differences in gene activity, based on gene connectivity and expand on the overarching sex-biased pathways. Next, we analyzed gene activity patterns and their regulation by sex down to the tissue level. Lastly, we provide examples of the importance of sex within these biological networks and why sex should be treated as a biological variable in systems biology.

## Materials and methods

### Study population

The Genotype-Tissue Expression (GTEx) project RNA-sequencing data (v8)^6^ was used to describe a whole-body map of sex differences in gene activity. Analyses were performed as shown in Figure 1 and Suppl. Fig. 1. To generate a whole-body map of sex differences in gene activity, we started our analysis with RNA-sequencing data from over 700 individuals in the GTEx project spanning more than 50 tissues. A comprehensive workflow of the analysis is shown in Fig. 1A (see Suppl. Fig. 1A for a more extensive flowchart). First, we selected samples that had at least a RIN > 6.0, a filter for the quality of RNA of the different samples. This cut-off is based on recommendations by the GTEx Portal. We performed principal component analyses on all tissues passing our criteria (Suppl. Fig. 2).

**Figure 1.**
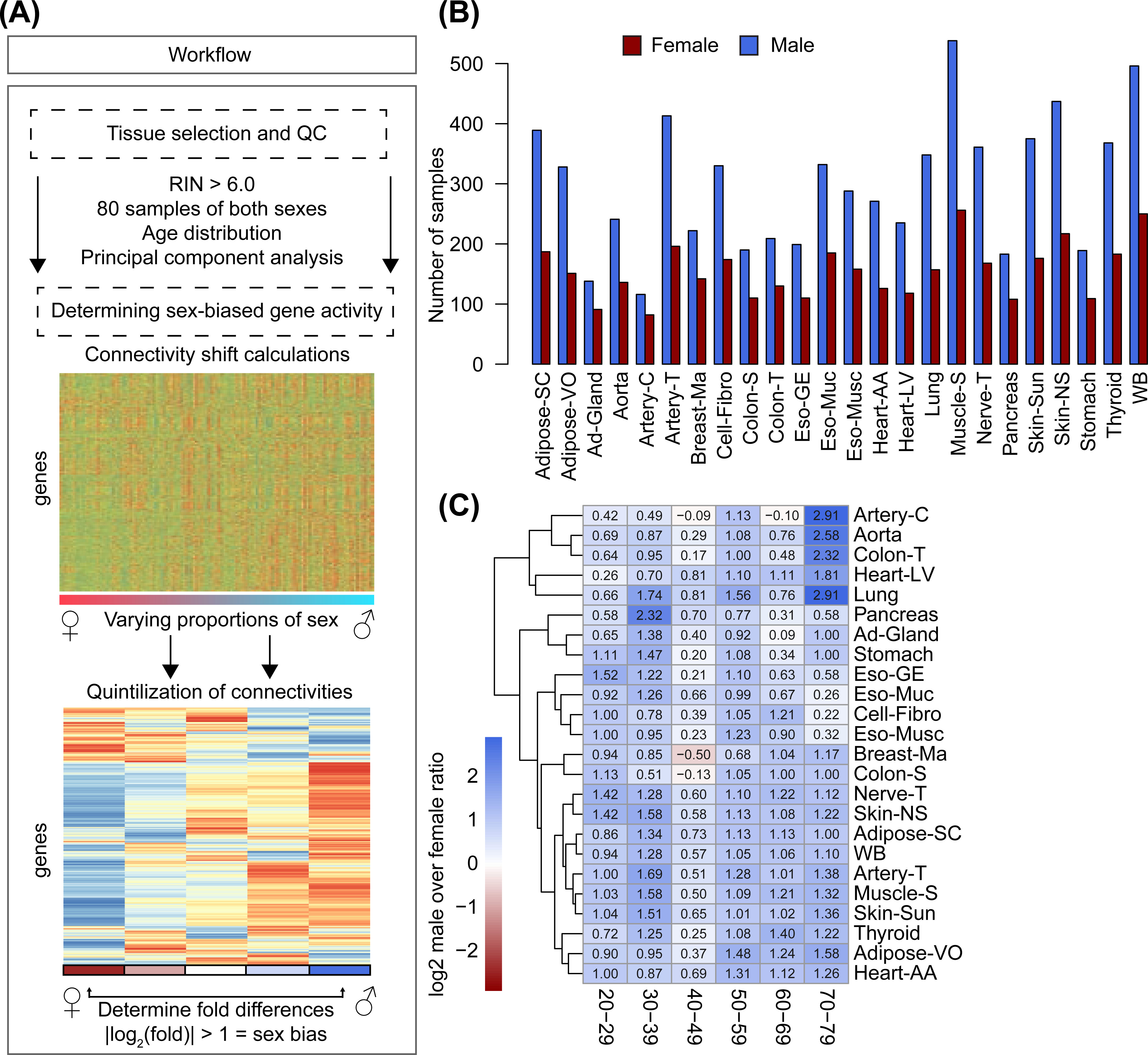
Workflow of sex-stratified gene activity analyses. A) The workflow of analyses performed in this study are shown. We started by analysing publicly available RNA-sequencing data from the GTEx Portal. We selected tissues with a RIN > 6.0, and with at least 80 samples of both sexes. Subsequently, we determined sex-biased gene activity in those 24 tissues that met our criteria. We calculated gene connectivities of genes over as much populations as 50% of the minimum sample size for one sex, but changing the population stepwise by sex, ranging from 100% females to 100% males. The left-most values indicate a population that is 100% female, while the right-most values indicate 100% male. Every step in between from left to right changes the ratio by removing one female sample and adding one male sample. Next, quintilization and cut-off by log-fold change differences selected genes that were sex-biased. See Suppl. Fig. 1A. for an in depth workflow. B) The amount of samples used over the 24 selected tissues is shown in a barplot. Red indicates female, blue indicates male. C) Log_2_ sex-ratios (male over female) are shown in a heatmap. The rows show the tissues, while the columns indicate age decades.

### Bioinformatics

Transcripts-per-million (TPM) expression data were downloaded from the GTEx Portal (v8), along with associated sample annotations and subject phenotypes. All analyses were performed in R (v3.5.1) within RStudio (v1.1.463). Principal component analyses and plots were done with 1000 genes that were most variable in expression for each tissue as defined by expression variance.

### Connectivity analyses

Genes for the whole-body map of gene activity were selected on the basis of variance and expression. A log_2_ expression cut-off of 1 TPM on average for each gene and variance of at least 1 were used to ensure enough information for connectivity analyses. Gene connectivity values were calculated using log_2_ TPM values for selected genes using WGCNA^7^ (v1.66) with the exponent equal to five. Repeated connectivity analyses using different populations with regards to sex-ratio are shown in Suppl. Fig. 1. Sex bias determination for individual genes is described in Suppl. Fig. 1. The cut-off for sex-bias was determined by permuting connectivity shifts 100 times using random samples in each population, instead of a gradient by sex (Suppl. Fig. 1A, box 3). We calculated log-fold changes of connectivity differences of 100 permutations in 3 different tissues; artery-C, heart-AA, and ad-Gland (Suppl. Fig. 1B). The amount of genes passing an absolute cut-off of log-fold change over 1 was always higher in connectivity shifts with a sex-gradient as compared to connectivity shifts with random samples, indicating that sex is an important determinant in connectivity (Suppl. Fig. 1C).

### Enrichment analyses

Hallmark enrichments were used to get an overview of well-curated lists of common pathways^8^. We defined sex bias in pathways by using the sex difference in number of genes for tissue-hallmark combinations. Those pathways with an absolute gene difference of at least 3 were deemed sex-biased (all plotted in Suppl. Fig. 4, and summarized in Fig. 2DE). E.g., if mammary tissue had 3 more genes that were male-biased for hypoxia than female-biased, mammary tissue would be male-biased for hypoxia. We set the cut-off at 3, as 95% of hallmark-tissue combinations had a lower absolute gene difference than 3.

**Figure 2.**
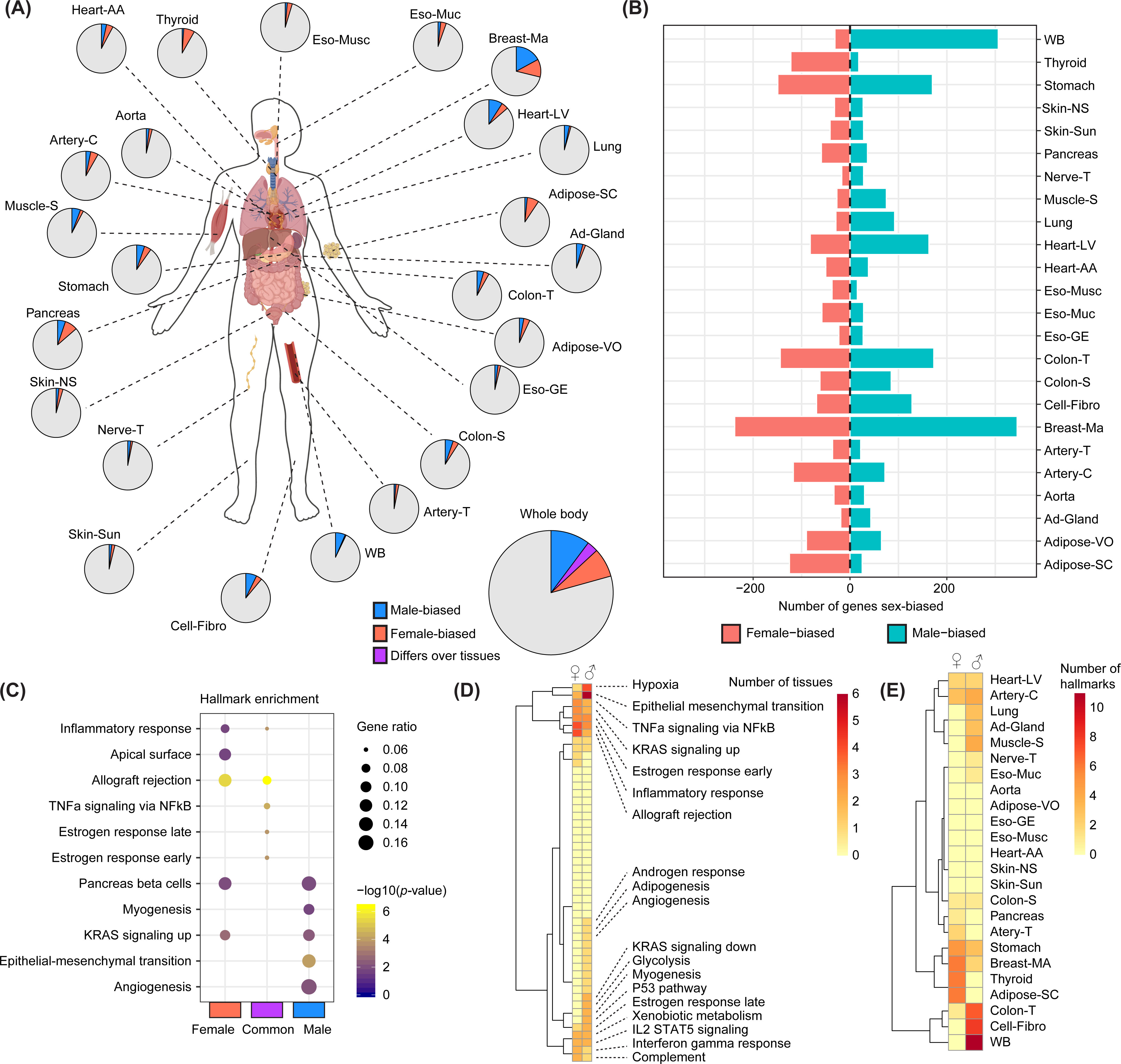
A whole-body map of sex differences in gene activity. A) A cartoon of the human body is shown with the tissues that are used in our study. For each tissue, the number of genes that are either male- (blue) or female-biased (red) are shown in a piechart. The bigger piechart shows all genes combined over all tissues, with purple indicating genes that differ in bias over multiple tissues. B) The exact number of genes that are either male- or female-biased per tissue is shown in a barplot. C) A dotplot is shown for the top hallmark enrichments for the whole-body genes that are female-, male- or either sex-biased. Color indicates *p-*value, while size of the dot indicates gene ratio. D) A heatmap is shown that indicates the number of times a certain hallmark enrichment with at least an absolute gene sex difference of 3 is either male- or female-biased. This is a summarized version of Suppl. Fig. 4A. A completely annotated heatmap can be found in Suppl. Fig. 4B. E) A heatmap is shown that indicates the number of times hallmark enrichments are either male- or female-biased over the different tissues. This is a tissue-level summarized version of Suppl. Fig. 4A.

### Motif analyses

Motif enrichment analysis was performed using RcisTarget^9^ (v1.2.1). In short, RcisTarget identified transcription factor binding motifs that were enriched in our lists of sex-biased genes. Motifs were subset to only the Hocomoco database. We set the cut-off for the normalized enrichment score (NES) to 0 to calculate NES for the majority of the motifs, after which we only took the ones calculated in both the female- and the male-biased genes (369 motifs). The motif rankings version used was human hg38 refseq-r80 for 500bp up and 100bp down the transcription start site. The motif-to-TF annotation version used was human v9.

### Permutations

We calculated enrichment for FDA-approved drug targets, targets for a male-biased disease (coronary artery disease (CAD)), and targets for a female-biased disease (multiple sclerosis (MS)) by permutation (10000x). FDA-approved drug targets were obtained from a comprehensive map of FDA drugs and their targets^10^. The CAD gene list has been described previously^11^, as well as a list for the major MS genome-wide association study^12^. Lists can be found in the Supplemental Dataset.

## Results

### The workflow of the study

The majority of samples within the GTEx Portal were male (Fig. 1B). Since we needed enough power to calculate sex differences in gene activity, we assessed tissues with at least 80 samples of both sexes. This reduced the number of tissues to 24, which were used in all analyses. We found that male and female samples were clustering mostly separately in breast-MA tissue, indicating that the transcriptome of male and female cells is most variable in breast-MA tissue. As the life expectancy of females is higher than males^13^, we determined the age distribution in male and female tissues. No major differences were found across the different life-decades in the tissues (Fig. 1C). Almost all the different age-tissue combinations were male-biased in sample size ranging from 1.06 times as many males in the adrenal gland (60-69 years of age) to 7.5 times as many males in the lung (70-79 years of age), indicating that the age distribution over all tissues is similar. One major exception was mammary breast tissue (breast-MA) in 40-49 years of age, which is female-biased (0.7 times as many males).

### A whole body-map of sex differences in gene activity

We found sex-dependent gene activity for 2,776 genes out of 13,787 (20.1%) genes tested over all tissues combined (Fig. 2A). The total number of genes that are exclusively female-biased only is 1,012, while 1,366 are exclusively male-biased. To generate a whole body-map of sex differences in gene activity, we calculated connectivity of highly variable and expressed genes in populations that differ in sex-ratio from 100% female to 100% male for every tissue (Suppl. Fig. 1A, box 1). Then, quintiles of populations varied by sex were formed by taking median connectivity values (Suppl. Fig. 3A). Sex-bias was subsequently calculated by dividing the connectivity value for the 5^th^ quintile (containing the majority of male populations) by the 1^st^ quintiles (containing the majority of female populations). Sex-dependent log-fold changes of all genes in all tissues can be found in Suppl. Fig. 3B. We noticed differences between the distribution of the gene connectivity of all tissues, e.g. activity in whole blood (WB) was more shifted towards males (Suppl. Fig. 3B). Of the 2,776 genes, 398 show sex-bias toward both sexes, but differed per tissue. The tissue that presented with most sex differences in gene connectivity is Breast-MA, with 345 genes male-biased and 238 genes female-biased. Tissues that presented mostly with male-activity were whole blood (WB) and skeletal muscle (muscle-S), while thyroid and subcutaneous adipose tissue (adipose-SC) presented mostly with female-activity (Fig. 2AB). Interestingly, fibroblasts, and thus cells *per se*, also presented with sex-bias in gene activity, indicating that tissue composition did not fully explain sex differences in gene activity. A complete list of genes that present with either male- or female-bias for each tissue can be found in the Supplemental Dataset.

### Consistent sex-dependent gene activity in pathways

Gene hallmark analysis showed that the entirety of female-biased genes over all tissues were enriched for immune response hallmarks, such as inflammatory response and allograft rejection (Fig. 2C). All male-biased genes together in all tissues were enriched for myogenesis, angiogenesis and epithelial to mesenchymal transition. The 398 genes that presented with either a male- or a female-bias over differing tissues were enriched for estrogen responses and inflammation pathways (Fig. 2C). To determine sex-dependent pathway activity over the different tissues separately, we calculated differences in gene presence of the different hallmark pathways (Fig. 2D, Suppl. Fig. 4A). Pathways that are more commonly male-biased over multiple tissues are hypoxia and epithelial to mesenchymal transition, whereas female-biased pathways are more inflammatory in nature. A fully annotated hallmark-heatmap can be found in Suppl. Fig. 4B. We also calculated which tissues are most commonly presenting with these sex-biased pathways (Fig. 2E). WB, fibroblasts, and colon transverse contain considerably more pathways that are male-biased, while subcutaneous adipose tissue (adipose-SC), breast-MA and thyroid contain more pathways that are female-biased.

### Multi-organ sex differences and regulation

As tissues in our body are connected by endocrine and paracrine mechanisms, we analysed gene activity patterns and their regulation by sex over multiple tissues together. For female-biased genes, 16.6% was female-biased in more than one tissue, while 12.7% of male-biased genes were male-biased in more than one tissue (Fig. 3A). The majority of these genes were consistently male- or female-biased in two tissues, as compared to more three and higher. An extreme example is the gene *USP32P2* which was female-biased in seven tissues, while the gene that was most consistently male-biased (in six tissues) was *SPATC1L*. To find exclusively male- and female-biased genes, we also removed genes that were male- or female-biased in different tissues. This led to 15.0% of all exclusively female-biased genes being female-biased in more than one tissue, while this number was 10.4% for male-bias. The most common exclusively male-biased genes were *PDK4, STC2, POMC, GSTM1, FCN3*, and *C4A*, being consistently male-biased in four tissues. The most common exclusively female-biased gene was *UQCRFS1P1*, being consistently female-biased in four tissues. Since the majority of genes, that exceeded singular tissue sex-bias, are sex-biased in two tissues, we looked at patterns in tissue-pairs in which these genes are biased (Fig. 3B). The most common female tissue-pairs were breast-MA with thyroid, adipose-VO with adipose-SC, and breast-MA with adipose-VO. This pattern was not present in male-biased genes, where the most common tissue-pairs were heart-AA with heart-LV, and breast-MA with colon-T. As differences in overall gene regulation may underlie consistent patterns of gene activity, we performed motif analysis on male- and female-biased genes. Normalized enrichment scores (NES) for 369 tested motifs differed between male- and female-biased genes, as visible from the differences in clustering (Fig. 3C). The top 10 motifs for both male-biased and female-biased genes are shown in Fig. 3D. Some potential transcription factors (TFs) driving higher gene activity for male-biased genes are *ZBTB18* and *TCF4/TCF12*. Some potential TFs for female-biased genes are *T, TBX21*, and the inflammatory regulators *RELA/NFKB1*, underlining the inflammatory nature of the female-bias.

**Figure 3.**
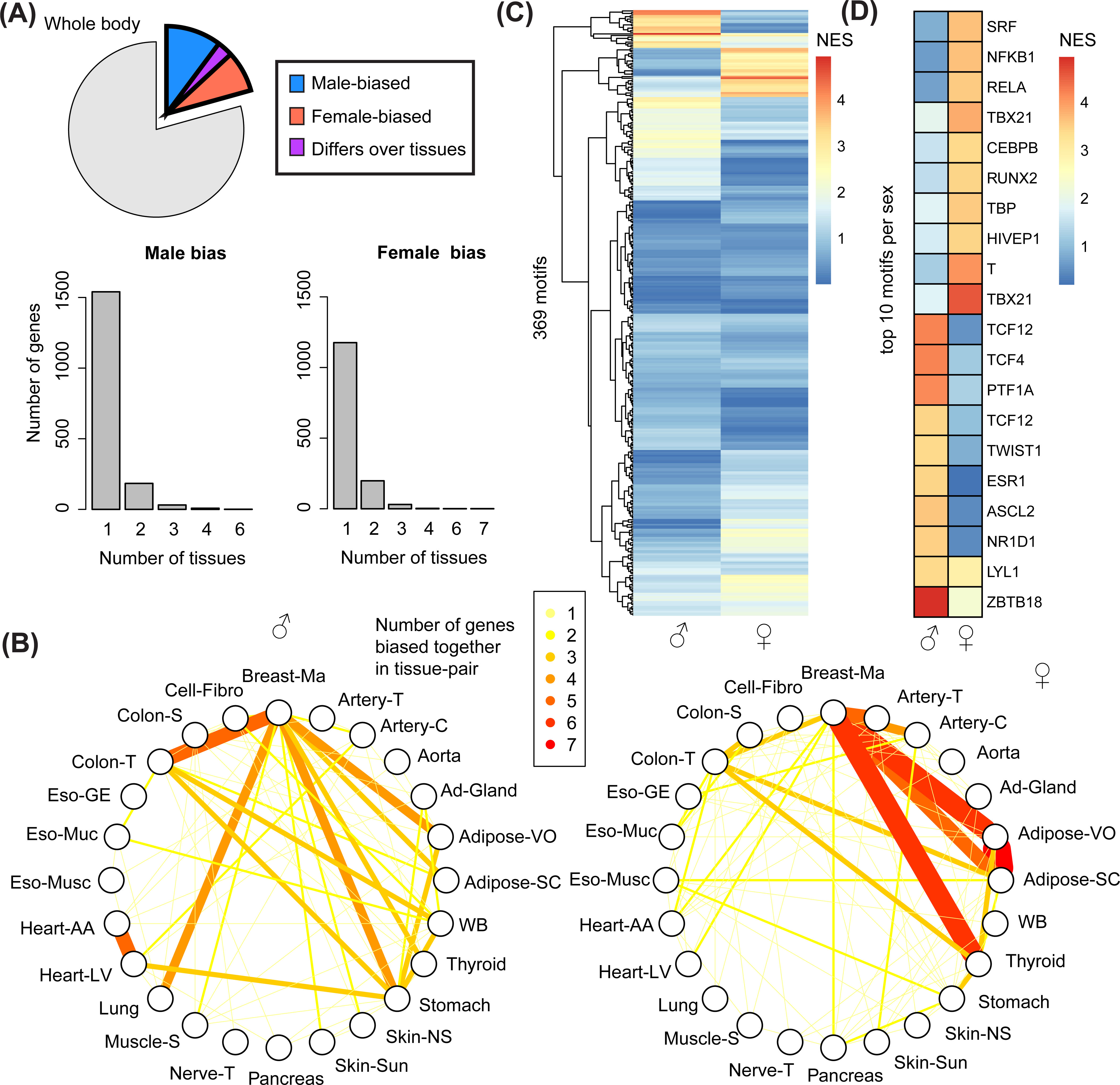
Tissue-transcending sex differences and regulation. A). A piechart indicates the percentage of genes with sex-bias collectively over the human body. The barplots show in how many tissues different genes are male- or female-biased. B) A graph plot shows the most common tissue-pairs for the genes that are male-biased (left) or female-biased (right) in two tissues. Color and edge weight indicate the number of genes that are biased in those two tissues. C) A heatmap shows the normalized enrichment score (NES) of 369 tested motifs for male-biased (left) and female-biased (right) genes in at least two tissues. A NES > 3 is considered significantly enriched. D) A heatmap shows the NES for the top 10 male- and female-biased motifs.

### Importance of tissue-specific sex dependent gene activity for target selection and drug discovery: Two examples

Systems biology provides an important step in the discovery of disease targets. To test the clinical importance of sex within systems biology, we calculated the overlap of sex-biased genes with 665 FDA-approved drug targets in the different tissues (Fig. 4A). The tissue with the most druggable sex-differentially active genes was mammary tissue, for both sexes (10 for males, 17 for females). Tissues with more male-active genes that were druggable targets were the adrenal gland, coronary artery, transverse colon, left ventricle, lung, and whole blood. Tissues with more female-active druggable targets were subcutaneous adipose tissue, visceral adipose tissue, mammary tissue, and thyroid. These results highlight the importance of tissue-specific sex dependent gene activity for drug targets for disease. Hence, within these lists, we highlighted a male-biased and a female-biased disease, coronary artery disease (CAD) and the autoimmune disease multiple sclerosis (MS), respectively (Fig. 4A). Within these sex-biased diseases, we identified sex-specific enrichment over different tissues as indicated in figure 4A for both CAD and MS.

**Figure 4.**
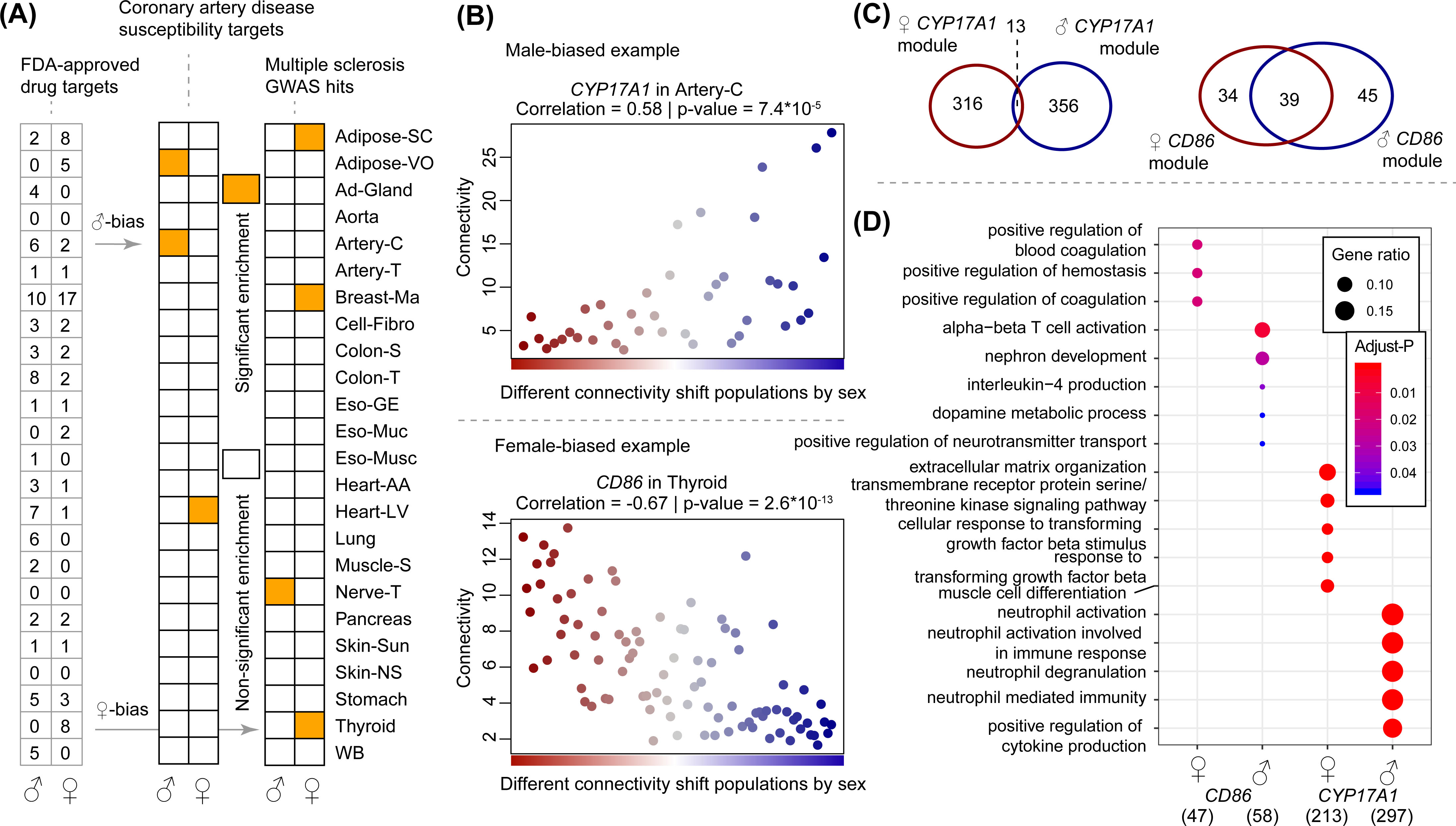
Use of sex-bias exemplified by FDA-approved drugs and sex-biased disease. A) A table shows the number of FDA-approved drug targets in the male- and female-biased genes (left). Two heatmaps shows whether or not male- or female-biased genes are enriched for coronary artery disease susceptibility targets and multiple sclerosis genome-wide association hits over the 24 different tissues. Orange indicates *p*_permutation_ < 0.05. B) Connectivity values for a male-biased (top) and female-biased (bottom) example gene that overlap with FDA-approved drug targets are shown over populations that differ in sex ratio. C) A venn-diagram shows the overlap of genes between sex-stratified Artery-C gene network modules that contain the *CYP17A1* gene (left), and the sex-stratified thyroid gene network modules that contain *CD86* (right). D) A dotplot shows gene ontology enrichment for biological processes in the sex-stratified *CD86* and *CYP17A1*-modules. Color indicates *p*-value, size of dot indicates gene ratio. The number beneath the sex-symbol indicates the number of annotatable genes of the module for that sex.

Following the analyses on tissue-specificity, we overlapped FDA-approved drug targets with sex-biased genes for both CAD and MS. For CAD, male-bias in coronary artery overlapped with the FDA-approved drug target *CYP17A1*, of which connectivity is higher in male populations as compared to female populations in artery-C (Fig. 4B, correlation between sex-ratio and connectivity = 0.58, *p* = 7.4e-0.5). *CYP17A1* is targeted by the FDA-approved drug abiraterone acetate, which is used to treat prostate cancer by lowering androgen levels^14^. This urged us to further construct sex-stratified gene networks in coronary artery. The male and female modules that contained *CYP17A1* were similar in size (369 and 329, respectively). The overlap, however, was minute, with 13 genes overlapping between the male- and female-*CYP17A1* modules (Fig. 4C). Gene enrichment analysis highlighted different biological processes in the male- and female-*CYP17A1* modules (Fig. 4D). The male module was enriched for neutrophil activation and cytokine production, whereas the female module was enriched for extracellular matrix organization and transforming growth factor beta responses. For MS targets, we overlapped FDA-approved drug targets with female-biased genes in thyroid tissue, since thyroid tissue is more often affected by autoimmunity in females as compared to males. The gene *CD86* was present in both gene lists and its connectivity was significantly correlated to sex-ratio (coefficient = −0.67, *p* = 2.6e-13). *CD86* is targeted by the FDA-approved drug abatacept for the treatment of rheumatoid arthritis, another autoimmune disease with higher prevalence in women^1^. Abatacept is more often discontinued in females as compared to males for the treatment of rheumatoid arthritis^15^. The sex-stratified gene modules in thyroid tissue for CD86 were similar in size, but did not materially overlap in terms of biological processes (Fig. 4C). The male module was enriched for alpha-beta T cell activation, while the female module was enriched for coagulation. These findings highlight the importance of incorporating sex in systems biology for improved understanding of diseases and search for novel drug targets.

## Discussion

We generated a whole-body map of sex differences in gene activity and highlight that sex is an important factor in systems biology: Overall, gene activity is sex-biased for 20.1% of the 13,787 genes analysed. As gene activity is the basis for further construction of regulatory gene networks to detect novel and druggable targets for disease, biological networks are more applicable to both women and men when analysed in a sex-stratified way. Interestingly, we found consistent sex differences in pathway activity present over multiple tissues in the human body. Furthermore, we showed gene activity varied by sex over multiple tissues, which was also accentuated by our analyses on regulatory DNA motifs. Lastly, we highlighted the importance of differences in tissue-specific gene activity by sex for target selection and drug discovery.

Sex-dependent gene activity might be driven by sex differences in (a combination of) sex chromosomes, sex steroids, and gender differences. The X- and Y-chromosomes harbor global gene regulators that are present in different doses between the sexes^16^. These genes can regulate the expression of genes located on the autosomes^17^, leading to differences in gene expression and activity between the sexes, and ultimately gene regulatory networks.

Sex steroids, such as estrogens and androgens, affect gene expression via multiple mechanisms. They may bind nuclear receptors, which subsequently translocate to the nucleus to bind promoter sequences, inducing or repressing transcription. Other mechanisms on transcription are less direct, such as interactions with epigenetic modifiers or signalling via G-protein coupled receptors. As the sex steroid arsenal varies immensely between males and females, sex steroids are strong candidates for influencing the sex-specific landscape of pathway activity. Importantly, we found estrogen responses to be enriched in genes that present with sex-bias in activity (Fig. 2CD). Remarkably, one of the genes exemplified in our study for CAD, *CYP17A1*, is implicated in the metabolism of sex steroids. In addition, the drug abiraterone acetate that targets *CYP17A1* is used in males to treat prostate cancer by lowering androgen levels^14^, highlighting a robust sex-specific use for targeting *CYP17A1*.

We found clear sex-dependent gene activity in general pathways over the human body. For example, pathways that were inflammatory in nature, such as TNF-a signalling and allograft rejection, were more commonly female-biased over multiple healthy tissues. This was also underlined by stronger enrichment for inflammatory and immune-related regulatory motifs in the promoters of these genes. Sex differences in the immune system have been thoroughly described^1^, and our data echoes the importance of sex in immunity. The tissue with the most female-biased (inflammatory) pathways was thyroid. Perhaps not coincidentally, autoimmune disease of the thyroid, such as Hashimoto’s thyroiditis and Graves’ disease, are more prevalent in females as compared to males^1^. Interestingly, MS-disease targets were also enriched in female-bias for thyroid tissue, but not in male thyroid. We found that the FDA-approved drug target *CD86* is female-biased in thyroid tissue. Strikingly, the drug that targets *CD86*, abatacept, has been discontinued more often in females as compared to males for the treatment of rheumatoid arthritis^15^, another female-biased autoimmune disease.

Enrichment for pathways such as myogenesis, angiogenesis and immune-related pathways may indicate differences in tissue composition between the sexes. We studied endothelial cell gene expression in GTEx as a marker for endothelial cell density (*VWF, CLDN5, CDH5 “https://gtexportal.org/home/gene/CDH5“*). The only tissue that showed a clear sex difference in endothelial cell expression was breast-MA. The strong sex difference in gene activity in breast-MA tissue may therefore also be driven by differences in cell composition, however, we cannot state this for other tissues, where endothelial cell gene expression did not differ as strongly. Interestingly, one of the strongest enriched transcription factor binding motifs (TFBM) for male-biased genes was *ZBTB18*. This gene has been shown to regulate the myogenesis genome network^18^. Myogenesis is one of the hallmarks that is overrepresented in the male-biased genes as well. *ZBTB18* might be sex-differentially affected to generate sex differences in body composition, such as for muscle mass^19^. One of the stronger TFBM for female-bias was *TBX21.* Mice lacking *TBX21* show increased visceral adiposity^20^, so an increased *TBX21* activity in females might help in explaining why females have less visceral adiposity as compared to males^19^. Tissue composition differences aside, we also found sex differences in gene activity and subsequent pathways in fibroblasts, highlighting that cells *per se* have different gene activity when comparing male to female cells which may contribute to sex differences in tissue composition.

### Limitations

An important message that needs to be reiterated is the underrepresentation of female samples in large databases, such as in our analysis (Fig. 1B). Unfortunately, this led us to exclude male samples from our analysis, to keep our studies equally powered for both sexes. We could not detect gender differences in our study, since the GTEx portal does not contain large questionnaires into behaviour and lifestyle choices of each individual. Larger epidemiological studies coupled to molecular deep phenotyping should allow for dissecting and understanding sex and gender influences simultaneously. Samples taken from individuals in GTEx are not specific for any disease and most likely represent healthy tissue. Sex-bias in specific disease settings might be different from those present in a healthy situation. Lastly, our analysis uses RNA-sequencing data, rendering us unable to look at protein levels and their modifications by sex, or at sex in the epigenetic landscape.

### Concluding remarks

The human whole-body map of sex differences in gene activity revealed major and consistent sex-dependency, in cells, single tissues, as well as over multiple tissues together. As gene activity is the basis for further construction of regulatory gene networks to detect novel and druggable targets for disease, biological networks should be analysed and interpreted in a sex-stratified way. This will accelerate drug development and ultimately benefit the health of both women and men.

## Abbreviations

Adipose-SC: Subcutaneous adipose tissue
Adipose-VO: Adipose tissue from the visceral omentum
Ad-Gland: Adrenal gland
Artery-C: Coronary artery
Artery-T: Tibial artery
Breast-MA: Breast mammary tissue
Cell-Fibro: Fibroblasts
Colon-S: Colon sigmoid
Colon-T: Colon transverse
Eso-GE: Gastroesophageal junction
Eso-Muc: Esophagus mucosa
Eso-Musc: Esophagus muscularis
GTEx: Genotype-Tissue Expression
Heart-AA: Heart atrial appendage
Heart-LV: Heart left ventricle
Muscle-S: Skeletal muscle
Nerve-T: Tibial nerve
Skin-Sun: Skin (sun exposed)
Skin-NS: Skin (non-sun exposed, suprapubic)
TFBM: Transcription factor binding motif
WB: Whole blood

## Author contributions

R.J.G. H. designed the study; analyzed the data; drafted the manuscript. H.d.R. supervised the study. All authors have contributed to scientific input and critically revising the manuscript.

## Acknowledgements

The Genotype-Tissue Expression (GTEx) Project was supported by the Common Fund of the Office of the Director of the National Institutes of Health, and by NCI, NHGRI, NHLBI, NIDA, NIMH, and NINDS.

