## Supplementary material for "Sex dependent gene activity in the human body": Suppl. Fig.

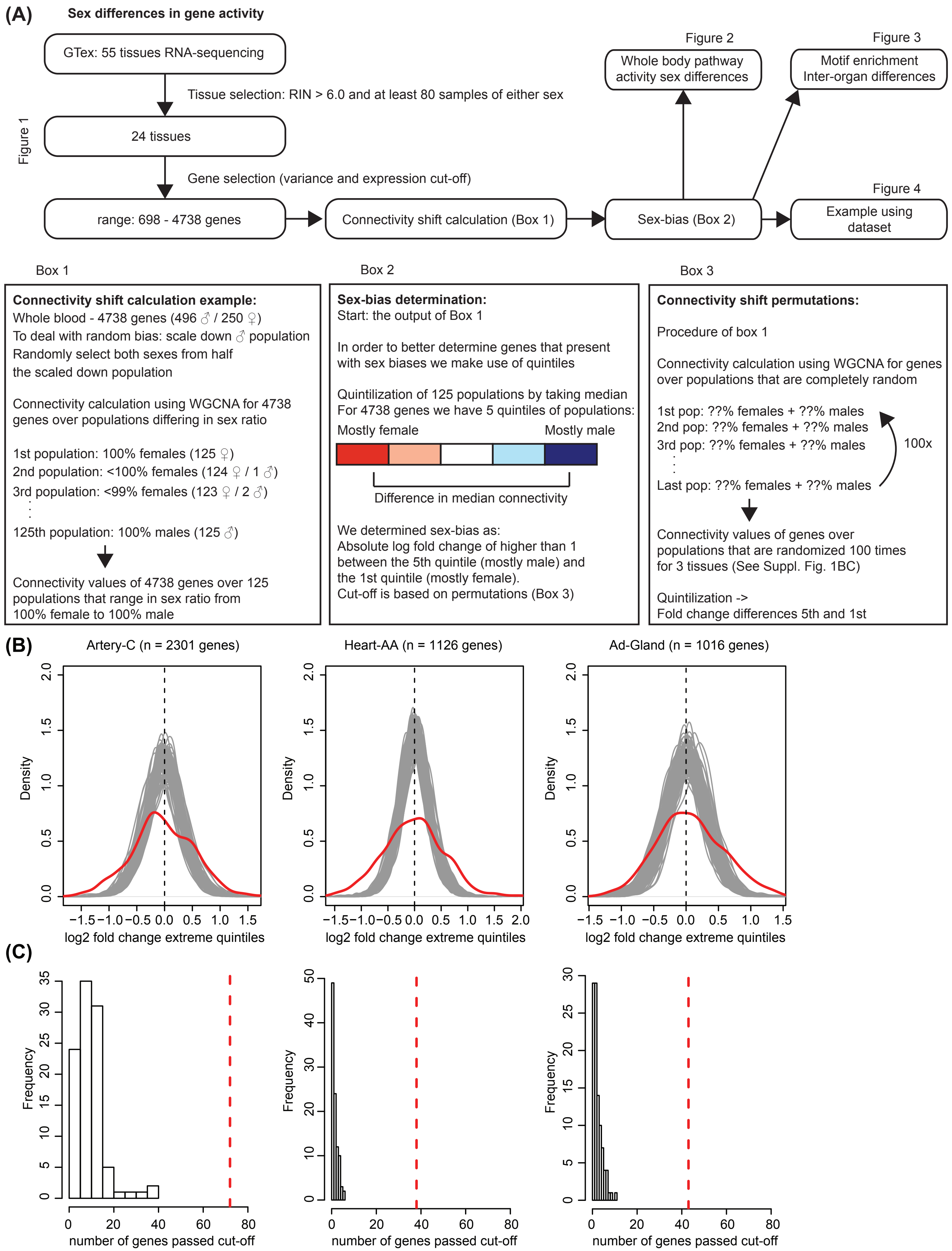


**Suppl. Fig. 1. In depth workflow analysis.** A) Workflow of the study and boxes to explain train of thought. B) Log fold change differences between the 5th and 1st quintile of the connectivity shifts are depicted for 100 permutations of random populations in artery-C, heart-AA, and ad-Gland (one grey = one permutation). The red line indicates the density distribution for the log fold changes of all genes in a sex-biased connectivity shift. C) Number of genes for the 100 permutatation cut-off log fold 1. The number of genes biased by a log fold 1 cut-off for the 100 permutations for 3 different tissues is shown in a histogram. The red dotted line indicates the number of genes with a log fold 1 cut-off in sex-biased connectivity analyses as used for the body map.

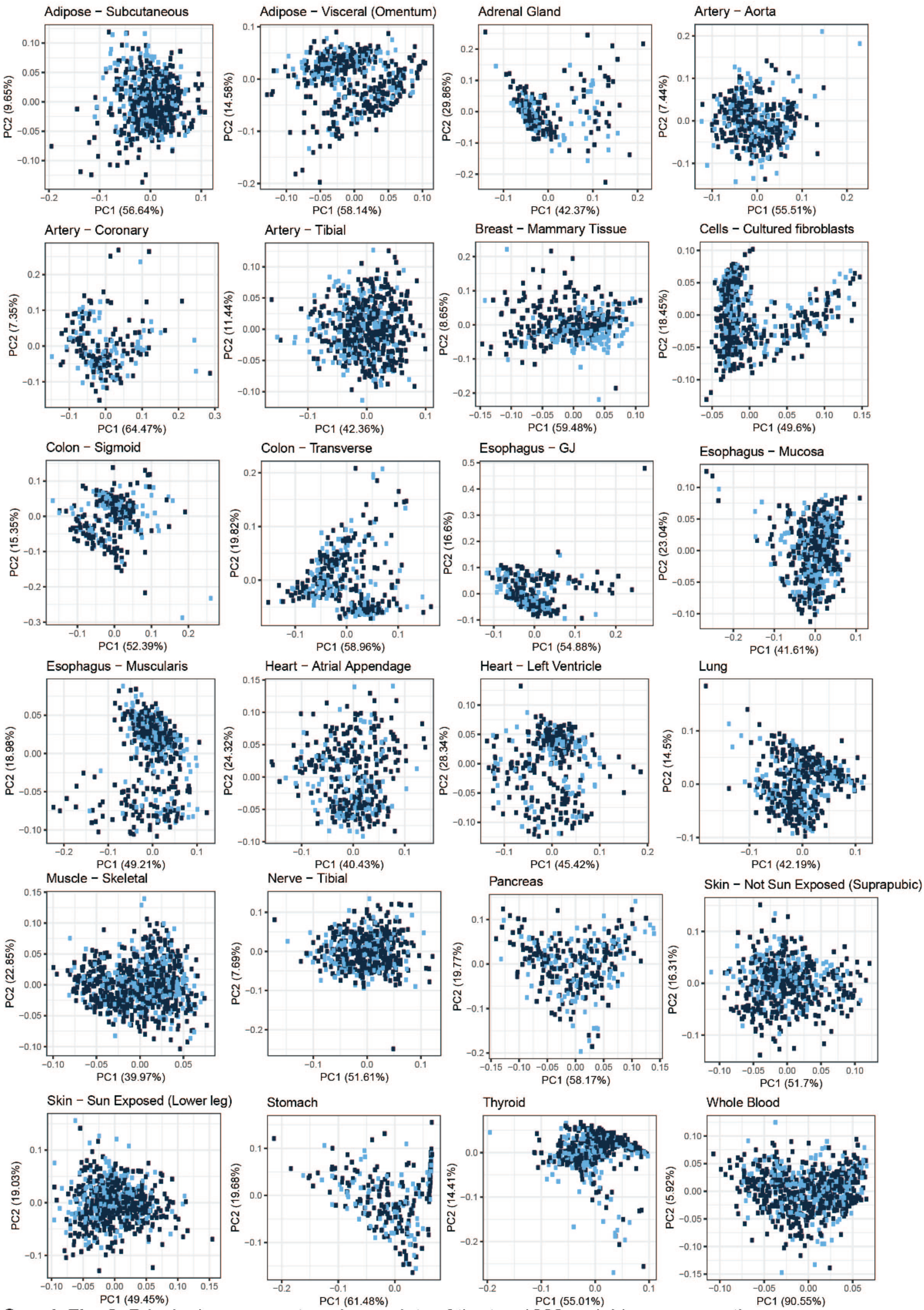


**Suppl. Fig. 2.** Principal component analyses plots of the top 1000 variable genes per tissue.
Lightblue = female, darkblue = male.

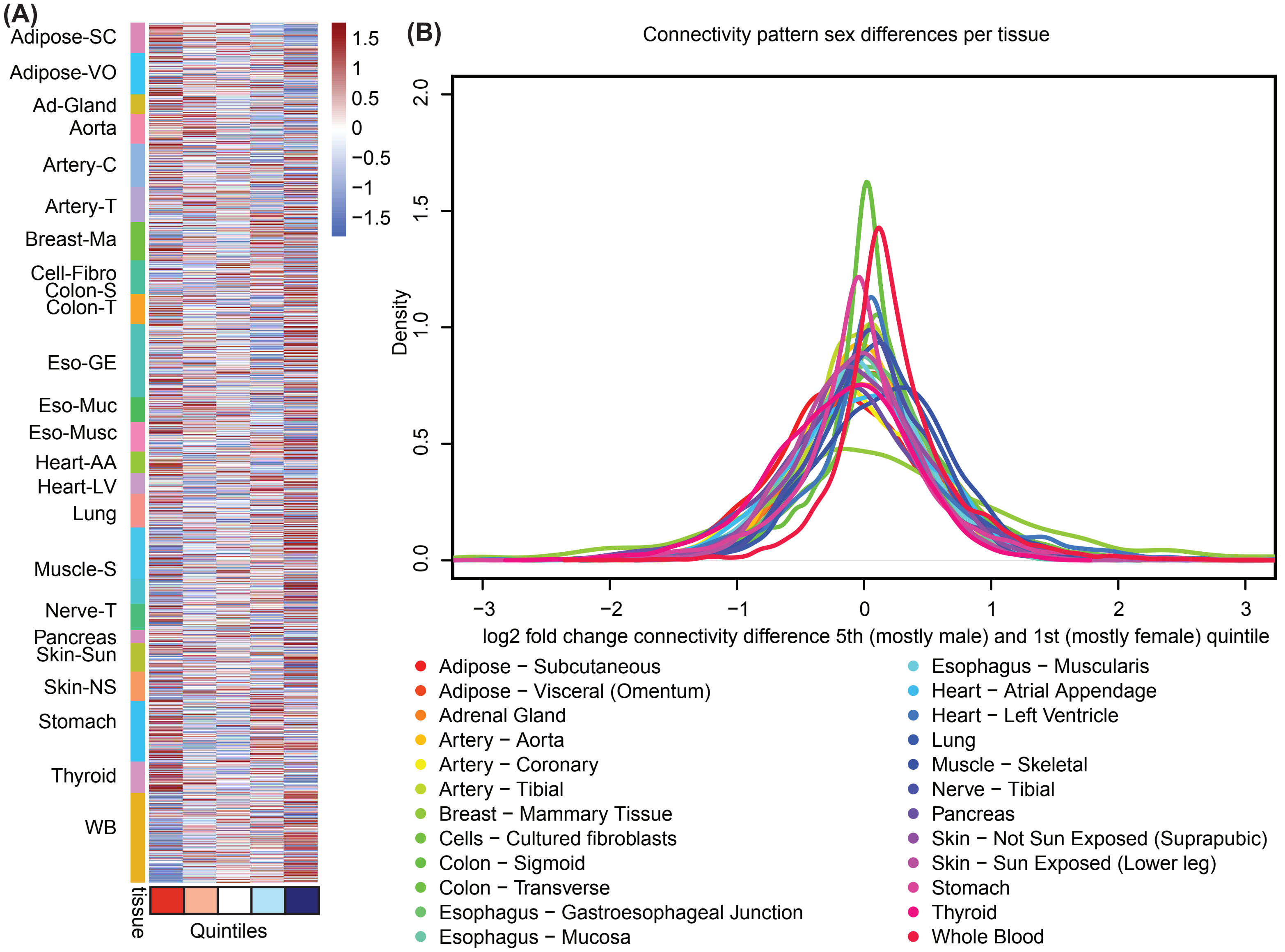


**Suppl. Fig. 3. Sex-bias in gene connectivity.** A) All connectivity values for all genes over all tissues tested are shown in a heatmap, divided in quintiles using their median. Quintile 1 = mostly female, quintile 5 = mostly male. B) Density plot for connectivity pattern differences. Log fold change differences between the 5th (mostly male) and 1st quintile (mostly female) of the connectivity shifts are depicted for all 24 tissues. Color indicates tissue.

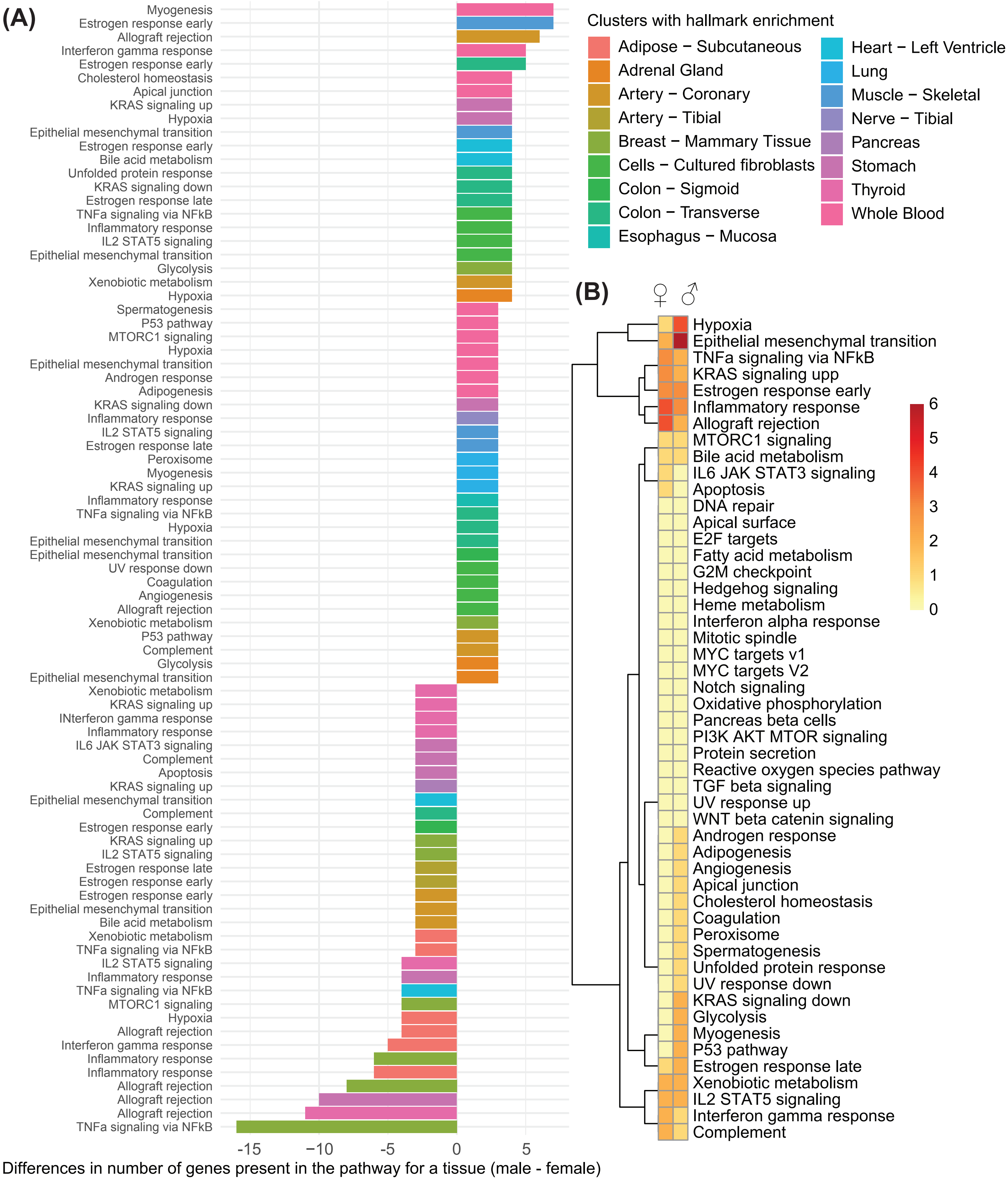


**Suppl. Fig. 4. Hallmark enrichment and sex-bias.** A) A barplot indicates the sex difference in number of genes for a tissue-hallmark combination that has at least an absolute gene difference of 3. Color indicates tissue, hallmarks are annotated on the left side. B) A heatmap is shown that indicates the number of times a certain hallmark enrichment with at least an absolute sex difference of 3 in gene presence is either male- or female-biased.
